# Structural and functional analyses of SARS-CoV-2 Nsp3 and its specific interactions with the 5’ UTR of the viral genome

**DOI:** 10.1101/2024.05.09.593331

**Authors:** Sofia Lemak, Tatiana Skarina, Robert Flick, Deepak T. Patel, Peter J. Stogios, Alexei Savchenko

**Author notes:** to whom correspondence should be addressed: Peter J Stogios,; or Alexei Savchenko.

## Abstract

Non-structural protein 3 (Nsp3) is the largest open reading frame encoded in the SARS-CoV-2 genome, essential for formation of double-membrane vesicles (DMV) wherein viral RNA replication occurs. We conducted an extensive structure-function analysis of Nsp3 and determined the crystal structures of the Ubiquitin-like 1 (Ubl1), Nucleic Acid Binding (NAB), β-coronavirus-Specific Marker (βSM) domains and a sub-region of the Y domain of this protein. We show that the Ubl1, ADP-ribose phosphatase (ADRP), human SARS Unique (HSUD), NAB, and Y domains of Nsp3 bind the 5’ UTR of the viral genome and that the Ubl1 and Y domains possess affinity for recognition of this region, suggesting high specificity. The Ubl1-Nucleocapsid (N) protein complex binds the 5’ UTR with greater affinity than the individual proteins alone. Our results suggest that multiple domains of Nsp3, particularly Ubl1 and Y, shepherd the 5’ UTR of viral genome during translocation through the DMV membrane, priming the Ubl1 domain to load the genome onto N protein.

## Introduction

The COVID-19 pandemic has brought into focus the danger and complexity of viral infections. As of the time of writing, SARS-CoV-2 caused more than 7 million deaths from more than 704 million infections (Worldometers.info). Research and development into direct-acting antivirals highlighted the necessity of detailed molecular understanding of the mechanisms of viral pathogenesis and the host-pathogen interactions that could be intercepted by antiviral therapeutics. In this vein, structural biology approaches have delivered stunning and rapid successes in providing molecular understanding into the SARS-CoV-2 proteins and their interactions with host factors (reviewed in (1, 2)). However, much remains to be learned about the molecular structure and function of some of this virus’ open reading frame products and their interactions with host proteins.

The SARS-CoV-2 virus’s genome represents a 30 kb single-stranded positive-sense RNA encapsulated by the Nucleocapsid (N) protein. The virion is protected by a host-derived membrane envelope harboring the Spike (S), Membrane (M) and Envelope (E) structural proteins. The SARS-CoV-2 genome encodes a total of 14 open reading frames which translate into 29 viral proteins. *orf1a* encodes the polyprotein pp1a, which is processed by the papain-like protease (PlPro) itself encoded within this open reading frame, into non-structural proteins (Nsps) 1 through 3. The genome also encodes *orf1b* and a -1 ribosomal frameshift upstream of the sequence corresponding to *orf1a’s* stop codon results in readthrough into Orf1b, which translates the polyprotein Orf1ab. Orf1ab encodes Nsps 4 through 10, which are liberated by the Main protease/Nsp5 region or Orf1ab.

Spanning 1945 residues the Nsp3 is the largest single protein encoded by the virus (reviewed in (3)). Along with Nsp4 and Nsp6, Nsp3 mediates the formation of double-membrane vesicles (DMVs) in infected cells, with the C-terminal portion (600 residues) of Nsp3 shown to be essential for this activity (4, 5). These virus-induced organelles are rich in double-stranded RNA. Accordingly, the DMVs are suggested to contain the viral replication-transcription (RTC) complex, shielding it from cytoplasmic RNA sensors that activate the innate immune system (6–10). Cryo-electron tomography (cryo-ET) studies of cells infected with SARS-CoV-2 or murine hepatitis virus (MHV) show that Nsp3 localizes to molecular pore structures embedded in the DMV membrane (9–12). Since the N protein is localized in the cytoplasm of infected cells (10), nascent viral ssRNA genomes must exit the DMV for packaging into a ribonucleoprotein complex with N. Similarly, viral mRNAs transcribed from the genome within DMVs must exist this organelle for translation by cytoplasmic/ER-associated ribosomes. Therefore, the Nsp3-containing molecular pore is thought to provide a key gate between viral RNAs in the DMV lumen and the cytoplasm, which facilitates Nsp3-RNA and Nsp3-N protein interactions to facilitate RNA exit and packaging (11–20).

Nsp3 from SARS-CoV-2 is comprised of at least seven structural domains (**Figure 1a**). While PlPro encoded as part of this protein has been the focus of intensive research as an established target of antiviral therapies, little is known about the molecular structure and function of other Nsp3 domains. A comprehensive study of the RNA binding properties of SARS-CoV-2 Nsp3 throughout the domains of this protein has not been carried out.

**Figure 1.**
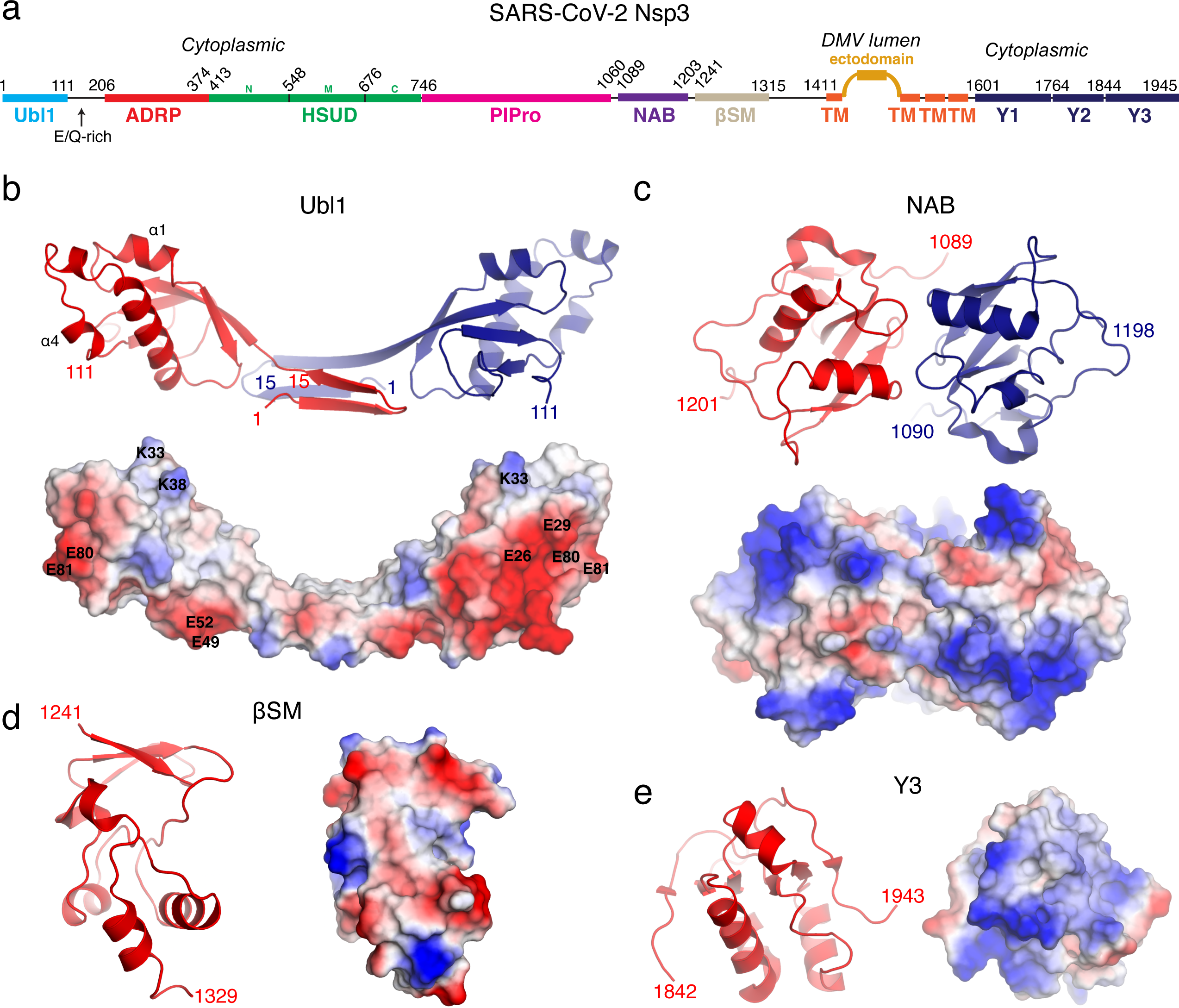

After PlPro, the next best-characterized region of Nsp3 is the ADP-ribose phosphatase (ADRP) domain (also known as the macrodomain or domain X). ADRP has been experimentally shown to harbor ADP-ribose phosphatase activity, has been the subject of numerous inhibitor screening campaigns (21–24) and has been shown to remove ADP-ribosyl groups from host PARP14 (25). Accordingly, this domain has been structurally characterised in complex with various ligands including small molecule inhibitors (24, 26, 27).

In contrast to ADRP, which appears to lack RNA binding activity, the Ubl1, human SARS unique (HSUD) and NAB domains of SARS-CoV Nsp3 have been shown to possess this activity (19, 20, 28–30). The Ubl1 domain of SARS-CoV Nsp3 was reported to have 20 µM affinity for ssRNA (20). It has also been predicted that the binding sequence for SARS-CoV Nsp3 Ubl1 is 5’-AUA-‘3, based on its co-purification with AUA-containing RNA, and biochemical characterization of short oligonucleotide binding (20). The HSUD domain of SARS-CoV and SARS-CoV-2 Nsp3 has been shown to bind G-quadruplex sequences (18, 28, 30, 31). The SARS-CoV Nsp3 NAB domain possesses nucleic acid-binding activity at micromolar concentrations to both ssDNA and RNA, with a preference for RNA substrates (32), and has been shown to bind to short, G-rich ssRNA, specifically those with three consecutive G residues (20).

A critical role of the Nsp3 Ubl1 domain in viral RNA synthesis has been attributed to its interaction with the N protein, which tethers Nsp3 to viral RNA during replication (16, 17, 33, 34). Genetic interaction assays have shown that the N-terminal region of MHV Nsp3, which contains Ubl1, binds to MHV N protein in an RNA-independent and species-specific manner (14, 35). In SARS-CoV-2, it has been shown that the Nsp3 Ubl1 domain interacts with the N-terminal domain of the N-protein as well as with two regions in the linker region between its N-terminal and C-terminal domains (34, 36). Since both proteins independently bind RNA and given the low occurrence of sequences predicted to be recognised by Ubl1 in the 5’-UTR or 3’-UTR, it has been speculated that the RNA-binding properties of Ubl1 have a role in its connection to N protein (33).

The final two domains of Nsp3 are the beta-coronavirus-specific marker (βSM) domain (also known as β2M) and the so-called Y domain which localises to the extreme C-terminus of Nsp3 (33). The Y domain has been subdivided into three regions: Y1, Y2 and Y3, showing significant variation across *Nidovirales*, and the Y2 and Y3 regions are restricted to *Coronaviridae* (32). The function of the Y domain of Nsp3 remains unknown.

In this study, we focused on expanding our understanding of the molecular features and function of the domains of SARS-CoV-2 Nsp3 outside of the PlPro domain, with a focus on interactions with viral RNA and the N protein. We determined the crystal structures of the Ubl1, NAB, βSM and Y3 domains of Nsp3, the last two of which provided the first molecular images of these regions of Nsp3. We show that five domains of SARS-CoV-2 Nsp3 interact with the 5’ UTR of the viral genome, including Ubl1, HSUD, ADRP, NAB and the Y domain. We demonstrate that the Ubl1-N protein complex shows higher affinity for the 5’ UTR than the isolated proteins, suggesting synergy between Ubl1’s N protein binding and RNA recognition. We also show that the Y domain also possess affinity to RNA, a function never attributed to this region of Nsp3. Altogether, these findings indicate that multiple regions of Nsp3 play important roles in shepherding the 5’ end of the viral genome through the DMV membrane for loading onto the N protein and suggest that they line the interior surface of the DMV pore.

## Results

### Crystal structures of the Nsp3 Ubl1, NAB, βSM and Y3 domains

As a first step in our functional analysis of Nsp3, we pursued structural characterisation of the individual domain in this protein. Ubl1 (residues 1-111), ADRP (residues 206-374), HSUD (residues 413-676), NAB (residues 1050-1216 and residues 1089-1203), the βSM (residues 1230-1334) were recombinantly expressed and purified from *E. coli*. Along the same lines, we expressed and purified the three fragments corresponding to Y1, Y2 and Y3 of Y domain (residues 1584-1945, 1619-1945 and 1844-1945) (**Figure 1a, Supplemental Data Fig. 1**). Using these purified Nsp3 fragments we were able to obtain structure determination quality crystals for Ubl1, NAB and the subregion Y3 region (residues 1844-1945) of Y domain (see material and methods for details). Notably, after depositions of the Ubl1 domain and the Y3 region structures to the publicly available database (PDB 7KAG, 7TI9 and 7RQG), the structures of the Ubl1-N protein complex and full-length Y domain were reported (34, 37).

The crystal structure of Ubl1 was solved by Molecular Replacement using the previously determined structure of the corresponding domain from the SARS-CoV virus (20). We determined the structure of this domain in two crystal forms (form 1 and form 2), both of which showed the same conformation of the Ubl1 protomer, suggestive of oligomerization architecture of the domain. In the case of the form 1 Ubl1 structure (**Figure 1b**) we were able to unambiguously assign all 111 residues of this fragment with two protomer chains present in the asymmetric unit. In contrast, the form 2 Ubl1 structure contained only one polypeptide chain in the asymmetric unit. However, the crystal symmetry of form 2 produced a dimeric structure identical to the dimeric structure in the asymmetric unit of form 1. Observing the same dimeric arrangement of Ubl1 fragments in two different crystal forms implied this to be functionally relevant for this domain of Nsp3. We observed the same oligomerization state for Ubl1 in size exclusion chromatography (**Supplemental Data Fig. 1b**). Dimerization interface in the crystal structures of Ubl1 was mediated by swapping of two N-terminal β-strands formed by residues 3 to 15. The crystal symmetry observed in the form 2 structure showed further association of two dimeric Ubl1’s into a tetramer, via extension of this N-terminal β-sheet (**Supplemental Data Fig. 2a**). To test the role of this region in oligomerization we designed and a purified Ubl1 fragment, missing fourteen N-terminal residues (Ubl1^Δ1-14^), and this remained monomeric based on size exclusion chromatography (**Supplemental Data Fig. 1b**). In line with significant sequence conservation between the SARS-CoV-2 Nsp3 Ubl1 domain with this domain in Nsp3 from SARS-CoV and MHV viruses (sharing 75% and 31% of sequence identity, correspondingly) the structures of these domains superimposed with RMSD of 2.7Å and 4.1 Å, over 102 and 100 C⍺ atoms, respectively vs Ubl1 from SARS-CoV-2, **Supplemental Data Fig. 2b**). This overall similarity is broken at the N-termini of the Ubl1 domains: this region in Nsp3s from SARS-CoV and MHV do not adopt the two β-strand arrangement observed in Ubl1 from SARS-CoV-2; they have been shown to be monomeric in solution (20, 35). This observation supports the notion that the N-terminus is the region responsible for dimerization of the Ubl1 domain. Our analysis of the electrostatic surface showed a clear acidic patch on one face of the Ubl1 from SARS-CoV-2, a neutral patch on the “top” face of the Ubl1, and the central domain-swapped region harbored largely neutral amino acids (**Figure 1b**).

The crystal structure of the NAB domain (**Figure 1c**) was solved by MR using the structure of the corresponding domain (residues 1089 to 1201) from SARS-CoV Nsp3 (19). We observed two polypeptide chains each corresponding to the NAR sequence in the asymmetric unit. However, the size exclusion chromatography showed that this domain is in a predominantly monomeric state in solution (**Supplementary Data Fig. 1d**). In contrast, we purified a 15-residue longer construct of the NAB, comprising residues 1050 to 1216, and this was dimeric in solution (**Supplementary Data Fig. 1e**), suggesting these additional 15 residues mediate dimerization. Further analysis of the NAB will focus on the dimeric structure evident in the crystal structure of the 1089-1201 construct, as this remains the only 3D structure available.

The determined structure of the NAB domain is highly similar to that of corresponding domain of Nsp3 from SARS-CoV (RMSD 0.9 Å over 113 matching Cɑ atoms). Importantly, the structures of NAB domains from these two viruses share common features in positioning of the residues K75/K74, K76/K75, K99/K98, and R106/R105 shown to contribute to RNA-binding (19). However, we also observed that the conformations of N- and C-termini differ between the structures of NAB from SARS-CoV and SARS-CoV-2 (**Supplemental Data Fig. 2c**). This difference may be due to these regions serving as flexible linkers with the PlPro and βSM domains and/or regions mediating dimerization.

The βSM domain does not share significant primary sequence similarity with any structurally characterised proteins. Therefore, we used the AlphaFold2 server (38) to generate a model of this domain and used it to solve βSM crystal structure by MR. The obtained structure is comprised of a three stranded β-sheet and short helices packing against the sheet spanning residues 1241 to 1329 of Nsp3 (**Figure 1d).** Notably, a search for structurally similar proteins to the βSM structure did not reveal any hits in the PDB database. The asymmetric unit contained 16 copies of the βSM domain. However, the size exclusion chromatography results (**Supplemental Data Fig. 1f**) showed this fragment to be predominantly monomeric in solution in line with PDBePISA server prediction of observed contacts between individual protomers in the crystal lattice which was not consistent with stable oligomerization. Our analysis of the βSM structure did not reveal any significant clefts or pockets that may be indicative of its molecular function.

While we were unable to obtain crystals of the full-length Y domain, we were successful with a fragment corresponding to its Y3 region. As in case of the βSM domain, the structure of Y3 fragment was determined by MR using an AlphaFold2-generated model. Retrospective analysis showed the AlphaFold2 model of the Y3 region closely matched its crystal structure with RMSD 0.4 Å over 84 matching Cɑ atoms (**Supplemental Data Fig. 3a**). Furthermore, the structure of the corresponding fragment in a consequently determined structure of full-length Y domain (PDB 8F2E (37)) also matched our Y3 region structure with a RMSD 0.5 Å over all 93 matching Cɑ atoms in this fragment. The Y3 crystal structure featured a mixed ɑ/β structure centered on a central 6-stranded anti-parallel β-sheet (**Fig. 1e**). Four chains were observed in the asymmetric unit of Y3 crystal lattice, with disulfide bounds formed via Cys1926 in each of two protomers pairs. However, both the Y3 fragment and full-length Y domain remained monomeric in solution according to size exclusion chromatography (**Supplemental Data Fig. 1g and 1h**), suggesting that the observed arrangements and covalent bonding between the protomers were a consequence of crystal packing and oxidation during crystallisation process, respectively. The Y3 fragment’s structure displays positively charged patches on its surface (**Fig. 1e**). A structural similarity search vs. the PDB showed that the Y3 domain show only very distantly similar matches (**Supplemental Data Fig. 2c)**; this lack of strongly structurally similar proteins suggested by our analysis is in agreement with that done using the structure of full-length Y domain (37).

### Multiple domains of Nsp3 bind the 5’-UTR of the SARS-CoV-2 genome in a specific manner

Previous studies have demonstrated that the domains of Nsp3 from MHV and SARS-CoV to bind ssRNA (20, 35). Thus, we explored the presence of such activity for domains of Nsp3 from the SARS-CoV-2 in a comprehensive manner. Based on previous work (20), we chose the 5’-UTR region of the SARS-CoV-2 genome as a potential substrate for Nsp3 domains. Electrophoretic mobility shift assay (EMSA) using [^32^P]-labelled ssRNA comprising bases 1 to 245 of the SARS-CoV-2 genome showed that the Ubl1 domain of Nsp3 binds this RNA fragment with Kd value of 31±3.9 µM, which is comparable to the affinity established for the corresponding domain from SARS-CoV Nsp3 (**Figure 2a**). To determine which charged residues were most significant for ssRNA binding, we used our crystal structure of Ubl1 to guide site-directed mutagenesis to alter negatively charged patches on the surface of this domain to positive charged and vice versa (**Figure 1b**, **Figure 2b**). The resulting Ubl1 variants were tested for RNA binding to the 5’-UTR substrate in comparison with the wild type Ubl1 (**Figure 2b**).

**Figure 2.**
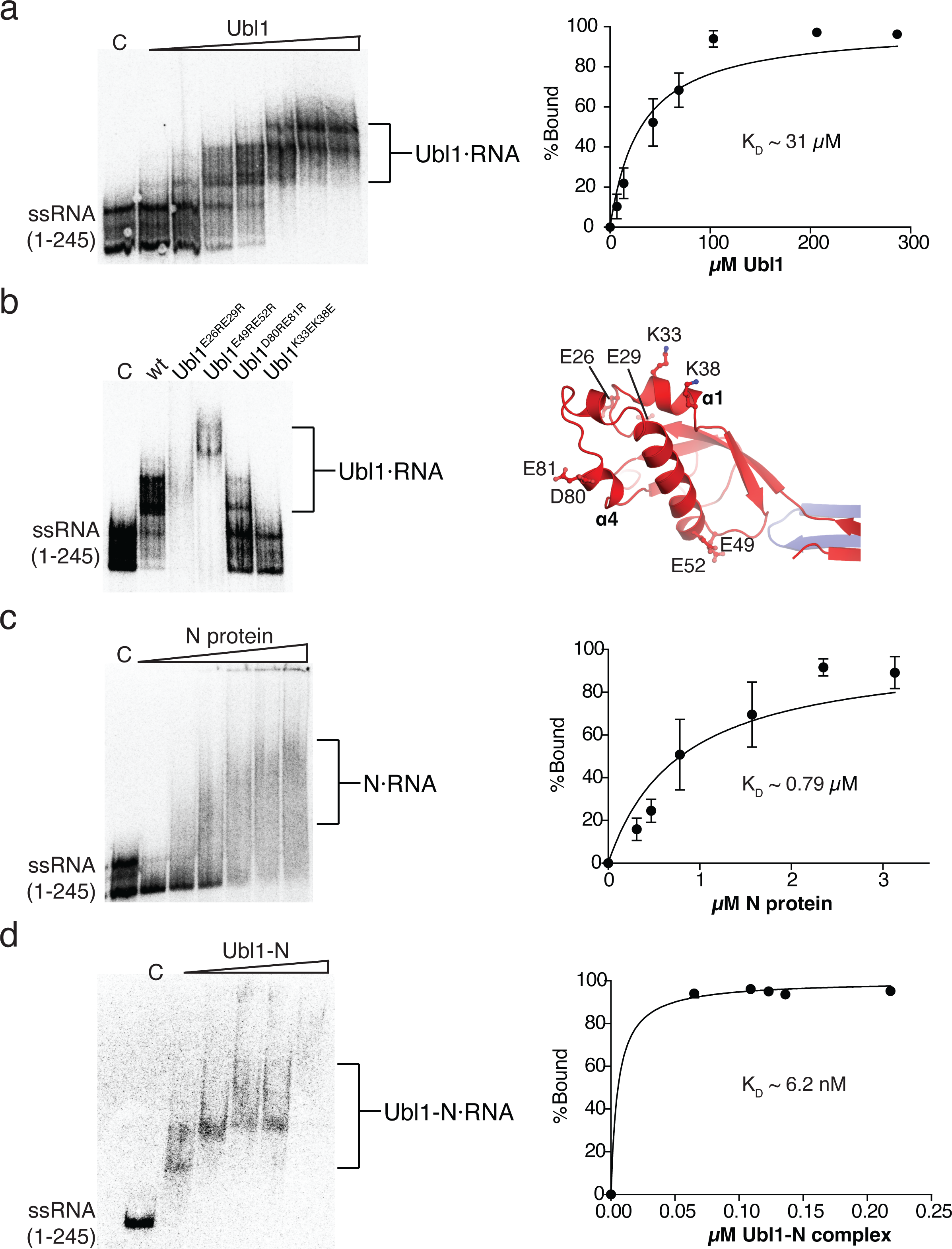

The Ubl1^K33EK38E^ variant carrying substitutions of residues located on the α1-helix, (**Figure 2b**) showed complete loss of ssRNA binding (**Figure 2b**). The Ubl1^D80RE81R^ variant with substituted negatively charged residues at the α4-helix to positively charged ones showed reduced affinity for ssRNA compared to the wild-type Ubl1. In contrast, the Ubl1^E26RE29R^ and Ubl1^E49RE52R^ variants showed ssRNA binding comparable to that of the wild type Ubl1. Interestingly, the Ubl1^Δ1-14^ deletion variant, which we showed was unable to dimerize, also showed complete loss of binding to 5’ UTR (**Supplemental Figure 3a**), while Ubl1^E26RE29R^, Ubl1^K33EK38E^, Ubl1^E49RE52R^ and Ubl1^D80RE81R^ all maintained the dimeric state adopted by the wild-type as verified by size exclusion chromatography (not shown) . This observation prompted us to suggest that dimerization is important for Ubl’s RNA binding.

EMSA analysis against 5’-UTR ssRNA 200 residue fragment showed that the ADRP, HSUD, NAB and Y (residues 1584-1945) domains of Nsp3 also show affinity to this substrate (**Figure 3)**. The calculated K_D_ values for these domains were 198±17, 41±4, 204±30, and 0.7±0.1 µM, respectively. These affinity values were lower than for Ubl1 with the notable exception of the Y domain, which demonstrated significantly stronger binding. Interestingly, the EMSA assay with the same substrate for Y3 fragment did not reveal any binding (**Supplemental Figure 3b**), suggesting the important role played in this activity the portion of Y domain corresponding to the Y1 and Y2 regions.

**Figure 3.**
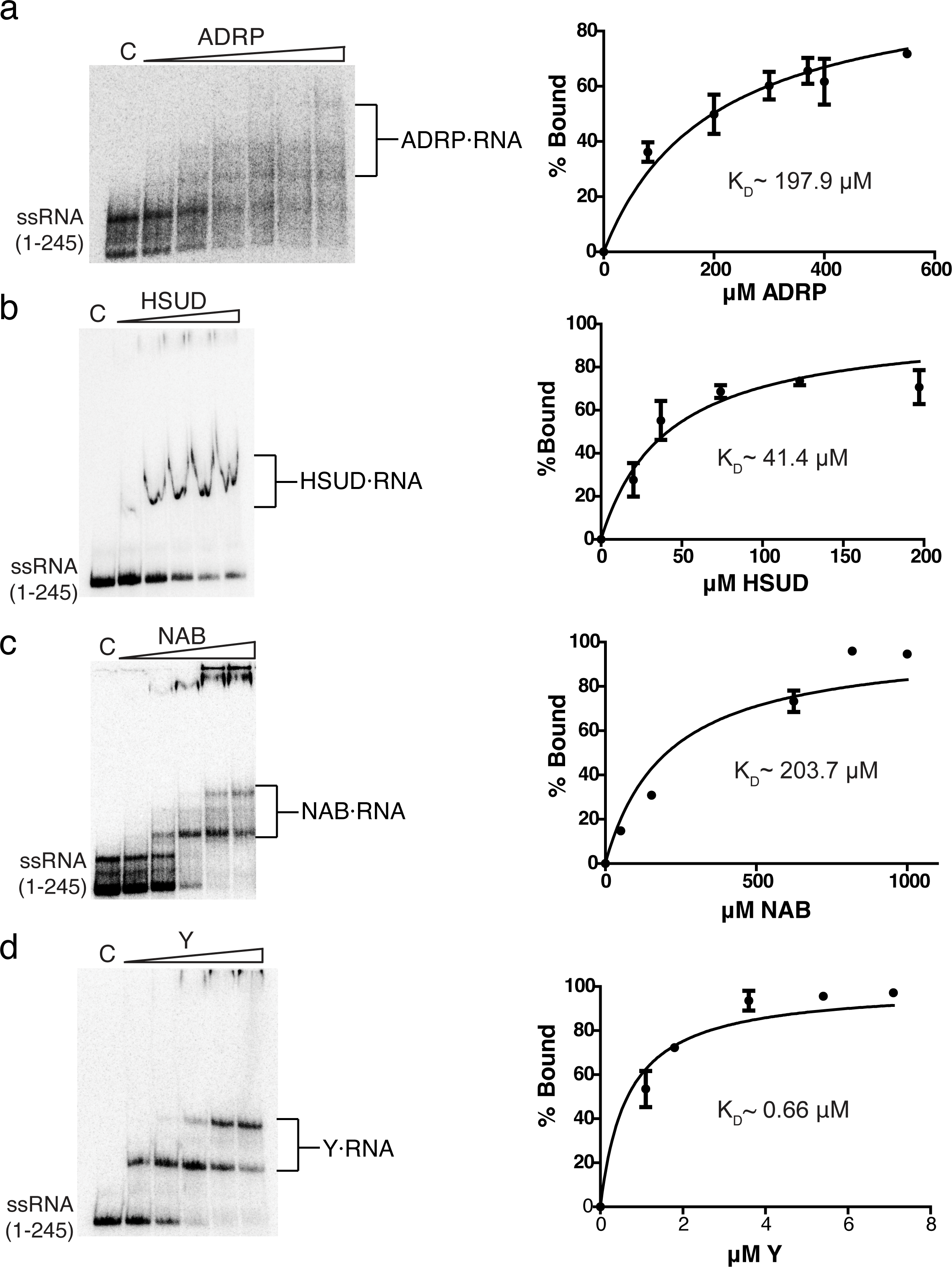

To characterize whether Nsp3 binds RNA in a sequence-specific manner, we tested whether the individual domains bind a region immediately downstream from the previously tested region of the 5’ UTR corresponding to 301 – 545 bases of the SARS-CoV-2 genome. We did not observe binding to this RNA substrate in cases of Ubl1, Ubl1^Δ1-14^, NAB, or Y domains of Nsp3. However, the HSUD domain did show binding to this RNA fragment (**Supplemental Figure 3c**). These results indicate that multiple Nsp3 domains specifically recognize the first 245 bases of the 5’ UTR, with HSUD possesses more promiscuous RNA binding activity.

### The Ubl1+N protein complex binds the 5’ UTR with higher affinity than the proteins alone

Since Ubl1 and N have been shown to form a complex involving the N-terminal domain and linker regions of the N protein (34, 36), we investigated how their interaction affected their binding to this RNA substrate using EMSA. According our results, the N protein binds to 5’ UTR with calculated K_D_ of 0.79±0.11 µM (**Figure 2c**). This value is higher than the calculated K_D_ (see above) for Ubl1 binding to the same substrate (**Figure 2a**). The EMSA assay against the same substrate using the His_6_-Ubl1 and N protein complex shows significantly higher affinity, reflected in calculated K_D_ of 6.2 ± 0.6 nM (**Figure 2d, Supplemental Figure 4**). This result shows that while both N protein and the Ubl1 domain of Nsp3 demonstrate significant affinity toward the 5’ UTR, binding is dramatically strengthened by formation of complex between these proteins. The Ubl1/N protein complex was also able to interact with 301-545 bp fragment downstream from the 5’ UTR (**Supplementary Figure 3c)**. This observation suggest that interactions with Ubl1 do not block the N protein from binding to this region of the viral genome; non-specific RNA recognition by N protein has been well established (39, 40).

## Discussion

The urgent necessity to develop therapies against SARS-CoV-2 infections have focused research efforts on individual proteins encoded by this virus. The analysis of Spike, RdRp and the two proteases PlPro and 3Cpro has been particularly intensive since these proteins represented the main targets of vaccination and antiviral therapies. These studies also highlighted the lack of extensive molecular knowledge about functional domains of Nsp3 protein beyond PlPro, even though it represents the largest non-structural protein encoded in the SARS-CoV-2 genome. To bridge this gap, we pursued the structural and functional analysis of multiple domains of this SARS-CoV2 protein, providing the molecular activities and the first experimentally derived structures in the case of the βSM and a region of the Y domain, which have never been experimentally visualised until this study.

We showed that multiple domains of Nsp3 including the Ubl1, ADRP, HSUD, NAB and Y domains recognize the 5’ UTR of the SARS-CoV-2 genome; this provides the first evidence of such activity in cases of ADRP and Y domains, and the first indication of the molecular function of the Y domain. Our mutagenesis analysis highlighted the role of individual surface residues in Ubl1 domain involved in interactions with viral RNA paving the way for further analysis of this activity. Future analysis based on our results will also be needed to localize the RNA binding surfaces on the other Nsp3 domains which we have demonstrated to possess such activity. In case of ADRP domain this activity will need to be reconciled with the enzymatic activity of this domain. Similarly, since the Y domain is expected to interact with Nsp4 and Nsp6, the effect of these interaction of RNA binding of this domain remains to be investigated.

Our analyses highlighted the role of Ubl1 domain of Nsp3 as the key connector between this protein, the N protein and the viral genome. Previous analysis using fluorescence polarization assay with short substrates (20 nucleotide) estimated the affinity of N protein to viral RNA to have the K_D_ of ∼7 nM (41). However, affinity decreased ∼10-fold when the protein was incubated with stem-loop (SL) RNA (41). This latter value is comparable with the one (0.79 µM) we obtained using EMSA assay against the 245 nucleotides of the 5’ UTR according for this protein. We observed a 127-fold increase in affinity for the N protein in the presence of Ubl1 domain as compared to N protein alone and 5000-fold as compared to calculated K_D_ for Ubl1 alone. This prompted us to suggest that the binding to RNA binding for Ubl1-N complex was synergistic, likely reflecting the *in vivo* importance of this complex formation during RNA exit from the DMV and its packaging onto N protein.

According to our assay the Ubl1 domain of Nsp3 shows specific binding to the first 245 bases of the 5’ UTR, in contrast to no binding to the sequence immediately downstream from this region. While there may be other part of the viral genome to which this domain has affinity, these results are supportive of Ubl1 possessing sequence specific RNA binding activity. Ubl1 specificity to the 5’ UTR is consistent with this domains established role in facilitating the binding of N protein to the first RNA bases exiting the DMV molecular pore and shielding it from cytoplasmic RNA sensors. Further studies will needed to delineate the specific sequence within the 5’ UTR recognised by Ubl1. Since the 1-245 fragment used in our assay includes predicted SL1 through SL4, and a portion of SL5, the role of these secondary structure elements (42) in interactions with Ubl1 should be also elucidated. Notably, the 5’ UTR contains three AUA sequences with one located in the 1-245 region, which were previously demonstrated to co-purify with SARS-CoV Ubl1, suggesting that these represent Ubl1 binding sites (20). Our mutagenesis and deletion analysis showed that the N-terminal β1 and β2 strands of Ubl1 as well as residues belonging to the α1 and α4 helices of this domain are important for recognition of the 5’ UTR. These results are consistent with previously reported mutational analysis of SARS-CoV Nsp3-Ubl1 complex which demonstrated that alteration of residues belonging to the α1 helix affected interactions with ssRNA (20). Another yet unclarified aspect of SARS-CoV2 virus’s life cycle is whether Ubl1 and the N-Ubl1 complex are able to discriminate between genomic RNA and subgenomic mRNA exiting the DMV pore. Presumably, the absence of the 5’ UTR on subgenomic mRNAs precludes their recognition by Ubl1, but further analysis is needed to clarify this.

During preparation of this manuscript, a study was published that delineated the interaction region between Ubl1 and the N protein and described the structure of this complex (36). The presented structure of the Ubl1-N complex contained only a single chain of Ubl1, while the sample used for structure reconstruction lacked the N-terminus (residues 1-15). A superposition of Ubl1 crystal structure with this domain in the complex with N protein showed that the conformation of N-termini (residues 1-14) does not introduce any steric clashes with the position of the N protein linker region, suggesting that dimerization of Ubl1 domain may be compatible with formation of complex with N protein. Observed interactions are also in line with analysis of Ubl1 equivalent domain called Nsp3a with N protein from MHV virus, where this interaction was mapped to ɑ2 of Nsp3a and the SR-rich region of N (35). The ɑ2 helices of SARS-CoV-2 Ubl1 and MHV Nsp3a are similar in structure (**Supplemental Data Fig. 2**) but show some primary sequence variation. Similarly, the N protein linker/serine-arginine (SR)-rich regions are similar between SARS-CoV-2 and MHV but do show variation in residue content. Specifically, the SARS-CoV-2 N protein contains one more arginine and two more serine residues, and one more hydrophobic residue in the hydrophobic region involved in interactions with Ubl1 according to our results.

The structure of SARS-CoV2 Ubl1 is very similar to that of the corresponding domains fromo SARS-CoV and MHV with exception of the N-terminus. In both crystal forms obtained for SARS-CoV2 Ubl1 we observed the fourteen N-terminal residues of this domain forming two β-strands involved in domain-swapping dimerisation. We further demonstrate that the deletion of this N-terminal portion of Ubl1 abolish both dimerization and the RNA binding functionality of this domain. Based on this analysis we hypothesised that the RNA binding surface and N-protein interaction surfaces of Ubl1 are fully formed only upon its dimerization. Further studies are necessary to define the role of Ubl1 dimerization in the context of full-length Nsp3 and its role in the DMV molecular pore.

We showed that the NAB of SARS-CoV-2 forms a stable dimer and possesses affinity toward 5’-UTR ssRNA substrates. Previous characterization of NAB from SARS-CoV, which was not reported to oligomerize, was shown to bind to A- and G-rich RNAs, such as (GGGA)_2_ and (GGGA)_5_ (19). The 1-245 and the 301-545 base 5’-UTR ssRNA substrates used in this study do not contain any GGG sequences, suggesting that NAB may recognise other RNA motives that remain to be characterised.

The HSUD domain of Nsp3 from SARS-CoV and SARS-CoV-2 share 75% of primary sequence identity. The HSUD of SARS-CoV-2 has been shown to bind G-quadruplexes/G4 sequences (18) and the HSUD from SARS-CoV has been shown to bind short RNA sequences generally rich in purines, as well as the TRS+ sequence in the 5’-UTR (29). While three PQS (Potential G4 Sequences) are predicted for the 301 – 545 base region, there are no PQS predicted for the first 245 nucleotides of the 5’-UTR of the SARS-CoV-2 genome (43–46). Since we observed binding of the SARS-CoV-2 HSUD to both these fragments of 5’-UTR, this suggests that this domain’s interactions with RNA may involve the sequences beyond PQS.

To our knowledge we are the first to report the RNA binding activity for the ADRP domain of SARS-CoV2 Nsp3. This domain adopts a compact structure featuring charged surface patches that can be responsible for observed RNA binding (24, 27, 47). However, the role of individual ADRP residues in interactions with RNA and how this activity relates to the catalytic and protein interaction activities reported for this domain ramains to be investigated.

The search for structurally similar proteins to the β2M domain did not reveal any significant hits suggesting that this domain adopts a unique fold. Since the β2M domain lacked any affinity to tested fragments of 5’-UTR the specific role of this domain remains unclear. Given its proximity to the transmembrane region of Nsp3, this suggests a potential role for β2M domain in orientation of the protein with respect to the DMV membrane and/or interactions with the membrane itself.

Overall, our results greatly expands the molecular data on individual domains of the largest protein encoded by SARS-CoV-2 virus. According to the current model SARS-CoV/CoV-2 viral genomes are shepherded onto the N protein by the Nsp3’s Ubl1 domain, which interacts with both the RNA itself and the N protein. The crystal structures of the Ubl1 and β2M domain presented in this study has been already used to validate the models of multidomain fragments or full-length Nsp3 obtained by cryogenic electron tomography (cryo-ET) (11, 12).

## Materials and Methods

### Cloning

The regions of SARS-CoV-2 *Orf1a* encoding the individual domains of Nsp3 were synthesized either by Twist Biosciences or using a BioXP 3200 (Codex DNA, San Diego, CA, USA) as codon-optimized for *E. coli* expression. As expressed as amino acids in mature Nsp3, domain boundaries of the individual domains were: Ubl1 1-111; ADRP 206-374; NAB 1089-1203 or 1050-1216; βSM 1230-1334; full Y 1584-1945; Y3 region 1844-1945. Synthetic dsDNA was then cloned into the pMCSG53 expression vector. Note that purified HSUD and PlPro were provided as gifts from the labs of Karla Satchell and Andrjez Joachimiak, respectively.

### Protein expression and purification

Expression plasmids were transformed into *E. coli* BL21 Gold (DE3) (Stratagene, San Diego, CA, USA) cells harboring an extra plasmid encoding three rare tRNAs (AGG and AGA for Arg, ATA for Ile) and proteins were overexpressed in 1 L in ZYP-5052 auto-inducing complex medium (48) by incubating a few hours at 37°C followed by transferring to 20°C for overnight growth. Cell pellets were collected by centrifugation at 6000 × *g*. Ni-NTA affinity chromatography was used for protein purification. Cells were resuspended in binding buffer [50 mM HEPES pH 7.5, 500 mM NaCl, 5% glycerol (v/v)], 0.5 mM Tris(2-carboxyethyl) phosphine (TCEP), 5 mM MgCl_2_, 1 mM phenylmethylsulfonyl fluoride (PMSF) and 1 mM benzamidine supplemented with 0.05% n-Dodecyl β-D-maltoside (DDM)] then lysed with a sonicator. After sonication and centrifugation (30 min at 20,000 rpm; Avanti J-25 centrifuge, Beckman Coulter, Brea, California, USA), cleared lysates were applied to nickel-nitrilotriacetic acid (Ni-NTA) resin. Beads were washed and proteins were eluted with loading buffer supplemented with 35 mM and 300 mM imidazole, respectively. Eluted His_6_-RNA-binding (residues 47-173) and His_6_-dimerization (residues 247-364) domains of the Nucleocapsid protein were dialyzed against 0.3 M NaCl, 10 mM HEPES pH 7.5, 2.5 mM MgCl_2_, 1 mM TCEP, 1% (v/v) glycerol. His_6_-N protein full-length alone and as a complex with His_6_-Nsp3 Ubl1 domain purified for pull down experiment and 4 mutants of Nsp3 Ubl1 (D80RE81R, E26RE29R, E49RE52R, and K33EK38E) purified for RNA binding assay were further purified by size-exclusion chromatography on a Superdex 200 HiLoad 16/60 column equilibrated with buffer composed of 0.5 M NaCl, 5% (v/v) glycerol, 10 mM HEPES (pH7.5), 2 mM MgCl_2_, 10 mM β-mercaptoethanol. Where necessary for crystallization, His_6_ tags were cut off by TEV protease (30 μg of TEV added to 1 mg of eluted protein) concurrently with dialysis at 4°C in either 300 mM NaCl, 10 mM HEPES pH 7.5, 1% (v/v) glycerol, 2.5 mM MgCl_2_, 1 mM TCEP (for Ubl1 and N protein), or 300 mM potassium chloride, 10 mM HEPES (pH 7.5) (for βSM) or 0.3 M potassium chloride, 0.5 mM TCEP, 2.5% (v/v) glycerol and 1 mM MgCl_2_ (for Y3 region). After dialyses, protein-TEV mixtures were passed through 2^nd^ Nickel-NTA to remove the His_6_ tags, TEV, and uncut protein. All proteins were concentrated using a BioMax concentrator (EMD Millipore, Burlington, MA, USA) followed by passage through a 0.2-μm Ultrafree-MC centrifugal filtration device (EMD Millipore, Burlington, MA, USA) and stored at -80 °C. Purity of proteins was checked using SDS-PAGE.

### Crystallization and x-ray structure determination

All crystals were grown at room temperature using the vapor diffusion sitting drop method using a Mosquito robot (SPT Labtech, Hertfordshire, UK). For Ubl1 form 1, 19 mg/mL protein was mixed with reservoir solution 1.6 M ammonium sulfate, 0.1 M HEPES pH 7.5 and 2% hexanediol and the crystal was cryoprotected with reservoir solution plus 30% ethylene glycol. For Ubl1 form 2, 10 mg/mL of the Ubl1-N protein complex was mixed with reservoir solution 1.6 ammonium sulfate, 0.1 M HEPES pH 7.5, 2% hexanediol and 1.25% 1-butyl-3-methylimidazolium dicynamide and the crystal was cryoprotected with reservoir resolution plus 25% ethylene glycol; note that only Ubl1 was found in the crystal. For NAB (residues 1089-1203), 15 mg/mL protein was mixed with reservoir solution 2 M ammonium sulfate, 2% hexanediol and the crystal was cryoprotected with paratone oil. For βSM, 15 mg/mL protein was mixed with reservoir solution 0.5 M MES pH 6, 40% tacsimate and the crystal was cryoprotected with paratone oil. For the Y3 region, 8 mg/mL protein was mixed with reservoir solution 1.1 M sodium citrate, 0.1 M HEPES pH 7.5 and the crystal was cryoprotected with paratone oil. Diffraction data at 100 K were collected at a home source Rigaku Micromax-007 rotating anode plus Rigaku R-AXIS IV detector, or, at beamline 19-ID of the Structural Biology Center at the Advanced Photon Source, Argonne National Laboratory. Diffraction data were processed using HKL3000 (49). Structures were solved by Molecular Replacement (MR) using Phenix.phaser (50) using the following models: for Ubl1, the Ubl1 domain from SARS-CoV (PDB 2GRI, (20); for NAB, the NAB domain from SARS-CoV (PDB 2K87, (19); for βSM and the Y3 region, models for MR were constructed by AlphaFold2 (38). Model building and refinement were performed using Phenix.refine and Coot (51). B-factors were refined as isotropic with TLS parameterization. Geometry was validated using Phenix.molprobity and the wwPDB validation server. Atomic coordinates have been deposited in the Protein Data Bank with accession codes 7KAG, 7TI9, 7LGO, 7T9W and 7RQG.

### Structural analysis

Oligomerization interfaces were analyzed using the PDBePISA server (52). Structural homologs in the PDB were searched for using the Dali-lite server (53) or the PDBeFold server (54). Electrostatic solvent-accessible surfaces were calculated using PyMOL (Schrödinger, LLC, New York, NY, USA). Figures were created using PyMOL.

### SEC-RALS of Ubl1-N protein complex

To clarify its molecular weight and suggested stoichiometry, the His_6_-Ubi1-N protein complex was produced by mixing individually purified His_6_-Ubi1 and N proteins, followed by size exclusion chromatography on a Superdex 200 HiLoad 16/60 column equilibrated with buffer composed of 0.5 M NaCl, 5% (v/v) glycerol, 10 mM HEPES (pH7.5), 2 mM MgCl_2_, 10 mM β-mercaptoethanol. Four peaks were observed in this chromatogram, with the second peak corresponding to the intact His_6_-Ubi1-N protein complex as indicated by SDS-PAGE. Further molecular weight and shape analysis of this peak containing the His_6_-Ubi1-N protein complex was carried out using size exclusion chromatography coupled with a 90° right-angle light scattering detector and 643 nm laser beam (OMNISEC Reveal, Malvern Panalytical, Malvern, UK). Before collecting any measurements, the protein was centrifuged at 10000 g for 30 min at 4 °C. The size exclusion analytical column (Bio-SEC-3, Agilent, Santa Clara, CA, USA) was loaded with 50-µl of protein at a concentration of 3.0 mg/ml. The protein was eluted through the column using a buffer composed of 250 mM NaCl, 20 mM HEPES pH 7.5, 5% glycerol, 5 mM MgCl_2_, and 10 mM TCEP. Analysis of the data was performed with the Malvern Analytical OMNISEC software. The molecular weight corresponded to 248,369, which approximately corresponding to a 4:4 complex.

### Preparation of nucleic acid substrates and electrophoretic mobility shift assay

The cDNA of SARS-CoV-2 was generated using the High Capacity cDNA Reverse Transcription Kit (Applied BioSystems, Waltham, MA, USA) from the MN908947.3 synthetic SARS-CoV-2 RNA (Twist Bioscience, South San Francisco, CA, USA). The DNA of the 5’-UTR region (1 – 245 bp) was amplified using PCR to include the T7 promoter with primers 5’-TAATACGACTCACTATAGGGATTAAAGGTTTATACCTTCC-3’ (forward) and 5’-GGACGAAACCTAGATGTGCTGATGATCG-3’ (reverse). The DNA of the region downstream of 5’-UTR (301 – 545 bp) was amplified using PCR to include the T7 promoter with primers 5’-TAATACGACTCACTATAGGG ACACGTCCAACTCAGTTTG -3’ (forward) and 5’-CTTCGAGTTCTGCTACCAGCTCAACCATAACATGAC -3’ (reverse).

Substrate ssRNA was transcribed using HiScribe T7 High Yield RNA Synthesis Kit (New England BioLabs, Ipswich, MA, SUA) and was [^32^P]-labelled at the 5’-end using T4 polynucleotide kinase (New England BioLabs) and purified as previously described (55). The reaction mixture for RNA binding assays with Ubl1, Ubl1 mutants, NAB, HSUD, and Y domains, as well as N protein and Nsp3-N protein complex contained 50 mM Tris-HCl (pH 8), 150 mM NaCl, 5 mM CaCl_2_, 1 mM DTT, 20 U RNaseOUT (Invitrogen, ThermoFisher, Waltham, MA, USA), and 8 nM (or 0.8 nM for reactions with the Nsp3-N protein complex) 5’-[^32^P]-labelled RNA substrate. Reaction mixtures for RNA binding assays with the ADRP domain contained 50 mM Tris-HCl (pH 8), 150 mM NaCl, 10 mM MgCl_2_, 1mM DTT, 20 U RNaseOUT (Invitrogen), and 8 nM 5’-[^32^P]-labelled RNA substrate. Reactions were incubated for one hour at 37°C, quenched by the addition of glycerol loading dye, and separated on 6% native polyacrylamide gels. Results were visualized using a Phosphoimager, with the percentage of bound substrate quantified using ImageLab software (Bio-Rad, Hercules, CA, USA). Values were plotted against total protein concentration to determine K_D_ values using non-linear regression fit in Prism software (GraphPad, San Diego, CA).

## Acknowledgements

We thank Rosa Di Leo and Cameron Semper for cloning. We thank the BioZone Mass Spectrometry facility for validation of protein purity. We thank Monica Rosas Lemus in Karla Satchell’s laboratory for purified HSUD protein, and Christine Tenar and Jurek Osipiuk in Andrzej Joachimiak’s laboratory for purified PlPro protein. We thank Changsoo Chang and Youngchang Kim at the Structural Biology Center, Advanced Photon Source, Argonne National Laboratory for x-ray diffraction data collection. This project was funded by the University of Toronto COVID-19 Action Initiative awarded to Aled Edwards. This project has also been funded in part with Federal funds from the National Institute of Allergy and Infectious Diseases, National Institutes of Health (NIH), Department of Health and Human Services, under Contract numbers HHSN272201200026C and HHSN272201700060C.

