## Supplemental Data for "Structural and functional analyses of SARS-CoV-2 Nsp3 and its specific interactions with the 5’ UTR of the viral genome"

### Supplemental Data Figure 1

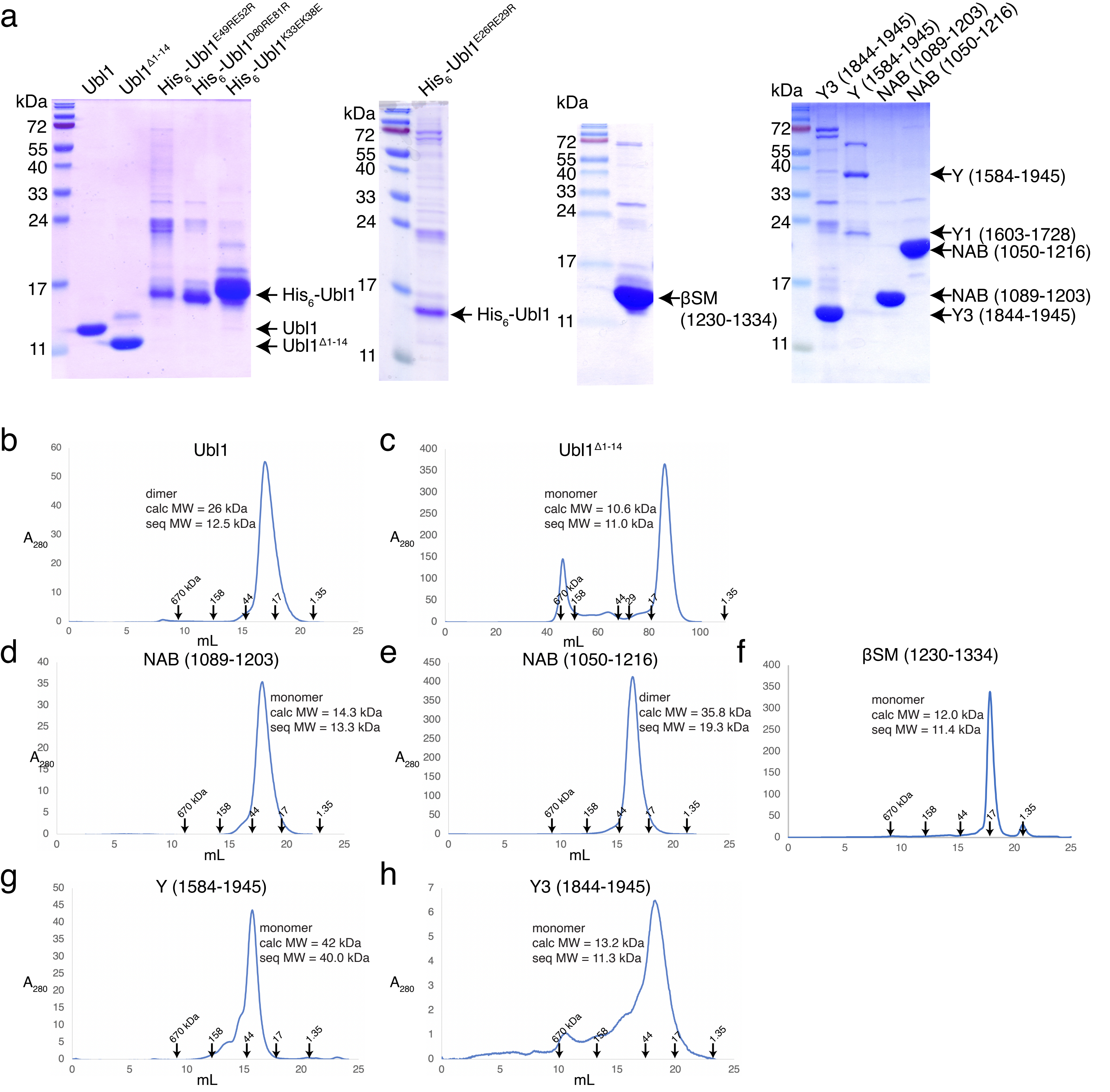

### Supplemental Data Figure 2

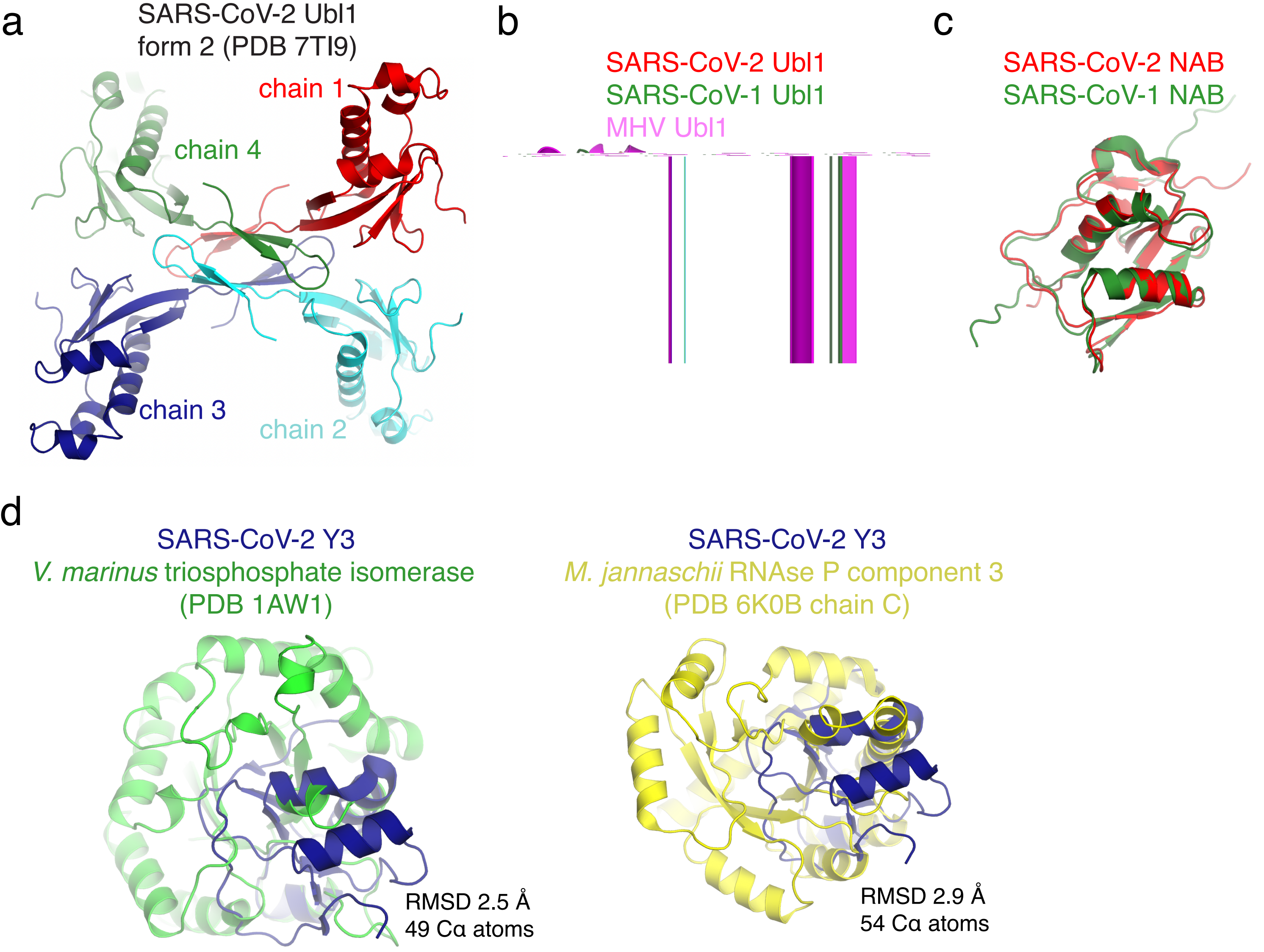

Supplemental Data Figure 3

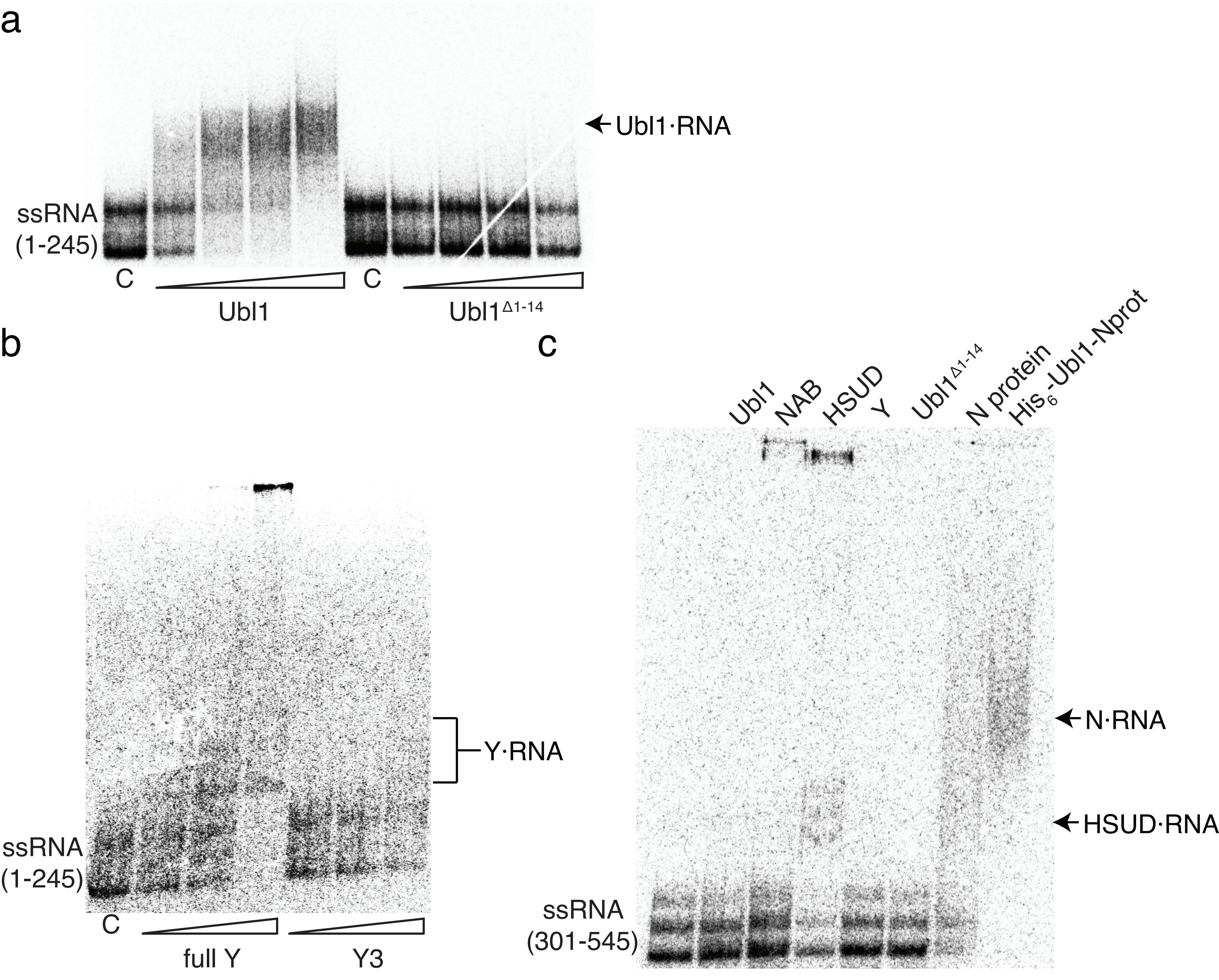

### Supplemental Data Figure 4

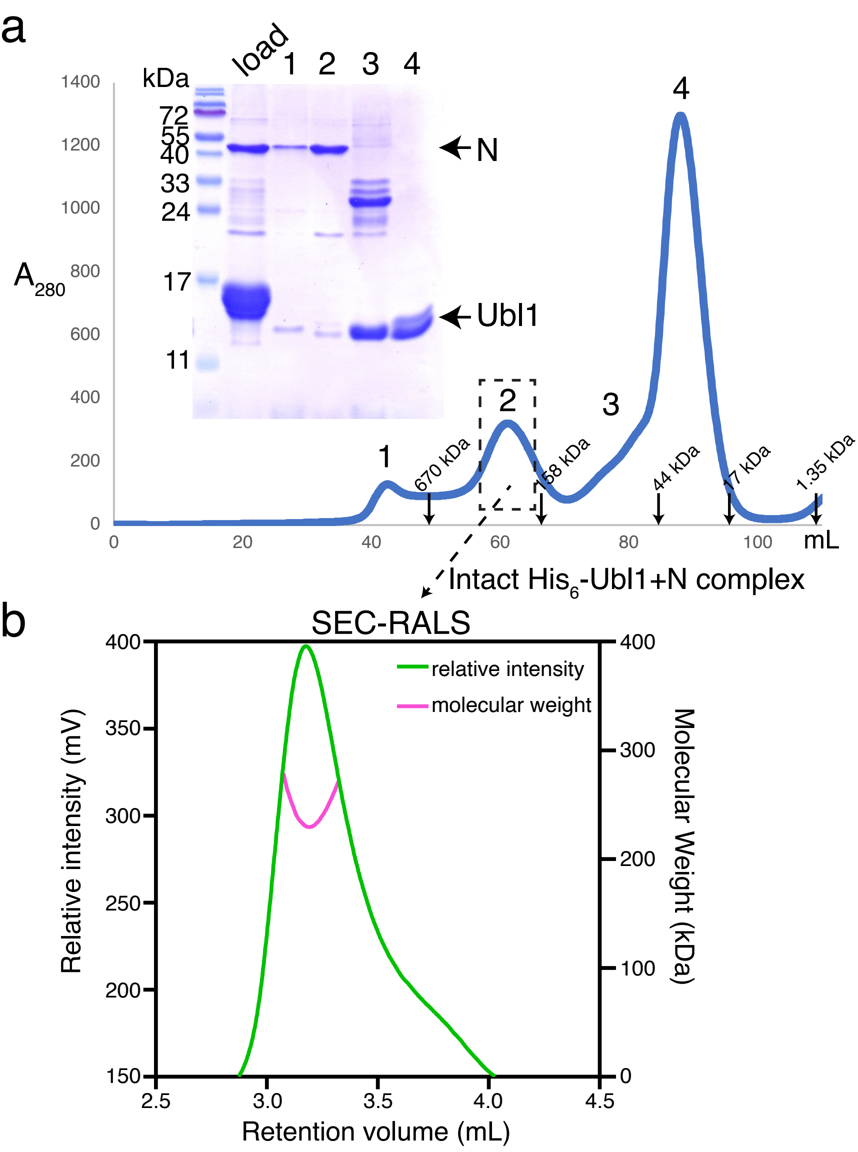


| Molecular weight (kDa) | Radius of hydration $R_h$ (nm) | Radius of gyration $R_g$ (nm) | Recovery of Sample (%) |
|---|---|---|---|
| 248,369 | 7.015 | 39.71 | 96.40 |
